# The UCSC Genome Browser database: 2016 update

**DOI:** 10.1101/027037

**Authors:** Matthew L. Speir, Ann S. Zweig, Kate R. Rosenbloom, Brian J. Raney, Benedict Paten, Parisa Nejad, Brian T. Lee, Katrina Learned, Donna Karolchik, Angie S. Hinrichs, Steve Heitner, Rachel A. Harte, Maximilian Haeussler, Luvina Guruvadoo, Pauline A. Fujita, Christopher Eisenhart, Mark Diekhans, Hiram Clawson, Jonathan Casper, Galt P. Barber, David Haussler, Robert M. Kuhn, W. James Kent

## Abstract

For the past 15 years, the UCSC Genome Browser (http://genome.ucsc.edu/) has served the international research community by offering an integrated platform for viewing and analyzing information from a large database of genome assemblies and their associated annotations. The UCSC Genome Browser has been under continuous development since its inception with new data sets and software features added frequently. Some release highlights of this year include new and updated genome browsers for various assemblies, including bonobo and zebrafish; new gene annotation sets; improvements to track and assembly hub support; and a new interactive tool, the “Data Integrator”, for intersecting data from multiple tracks. We have greatly expanded the data sets available on the most recent human assembly, hg38/GRCh38, to include updated gene prediction sets from GENCODE, more phenotype- and disease-associated variants from ClinVar and ClinGen, more genomic regulatory data, and a new multiple genome alignment.

## INTRODUCTION

The University of California Santa Cruz (UCSC) Genome Browser (1, 2) is a publicly available collection of tools for visualizing and analyzing both the large repository of data hosted at UCSC and user-supplied data. We aim to provide quick, convenient access to high quality data and tools of interest to those in the academic, scientific, and medical research communities.

The UCSC Genome Browser database hosts a large repository of genomes with 166 assemblies from GenBank (3) that represent over 93 different organisms across the tree of life, from vertebrates such as human, mouse, and zebrafish to insects and nematodes. Each of these assemblies has a standard set of basic annotations that includes assembly contig and scaffold names, gap locations, GC percent calculated in 5-base windows, gene predictions using Genscan (4), CpG islands (5), and repetitive elements identified using RepeatMasker (Smit, AFA, *et al., RepeatMasker Open-4.0* at http://www.repeatmasker.org), Tandem Repeat Finder (6), and WindowMasker (7). When the data are available, ESTs and mRNAs from GenBank and genes from RefSeq (8, 9) are mapped to the genome. Beyond these basic annotations, some assemblies offer additional data sets such as gene predictions from Ensembl (10, 11) or other sources, sequence alignments using Lastz (12) and Multiz (13), conservation scores based on phastCons (14) and phyloP (15), and chain/net (16) pairwise alignments with other assemblies. The richly annotated human and mouse genomes offer a wider range of data including gene predictions from GENCODE (17), regulatory and expression data from ENCODE (18, 19), phenotype and disease associations from various sources, variation data from dbSNP (20), and complex repeat structure visualization.

In addition to hosting and visualizing genomes and their associated annotations, the UCSC Genome Browser provides a suite of both web-based and command-line tools to assist researchers in analyzing these data. Some of these web-based tools offer alternate visualizations of a data set, such as Genome Graphs, while others facilitate searching of the genome, such as BLAT (21), or examination of the underlying data, such as the Table Browser (22) and the Data Integrator. Our command-line tools allow for the interrogation of large data files and conversion among numerous file types, and provide pipeline-ready versions of our popular web-based tools such as BLAT or LiftOver.

In the following sections, we first offer a brief overview of the Genome Browser’s major features, then highlight some of the substantial data sets and software improvements added in the last year, and conclude with hints at future directions.

## OVERVIEW OF THE BROWSER

### Visualizing and navigating data

The UCSC Genome Browser provides an interactive web-based platform for visualizing large genomic data sets (or “tracks”), including gene predictions, expression levels, and regulatory elements alongside one another. The user interface offers a broad range of customization of both the display of the available tracks as well as the browser window itself. The track display is dynamic, and tracks can be added, removed and rearranged in the browser display at will. The visibility mode of each track can be adjusted using different settings ranging from those that show detailed individual annotations to compact density graphs that summarize the data. The Genome Browser display also includes a number of options for navigating between regions of the genome. Users can quickly zoom to different levels of genomic resolution, from a single base pair to an entire chromosome, or scroll across the genome with the click of a button. It includes a search box that accepts genomic positions or search terms ranging from gene symbols to accession numbers. Users can click on individual items to see detailed information about the annotation and learn the methods used for generating it. After configuring the Genome Browser display, users can save the configuration to a “session” for later viewing or sharing with colleagues. Any custom data present when saving a session will also be preserved.

### Incorporating user-generated data

One of the most powerful features of the UCSC Genome Browser is the ability for users to upload and visualize their own data alongside UCSC-hosted tracks through custom tracks and data hubs. Small data sets can be uploaded and viewed in the browser as custom tracks using a variety of formats including BED, WIG, pgSnp, GFF, GTF, MAF, and PSL. Larger data sets and collections of data sets are better suited to organization and remote access through the track data hub mechanism (23). While track data hubs support a more limited set of file types (bigBed, bigWig, bigGenePred, BAM, HAL and VCF), they provide users with greater control over the organization, display, and sharing of their custom data sets. Hubs also allow datasets larger than the upload limits for custom tracks. The bigBed, bigGenePred and bigWig file formats (24) can be generated from their respective uncompressed file types using command-line tools available from UCSC (http://hgdownload.soe.ucsc.edu/admin/exe/). Assembly data hubs, which are an extension of track data hubs, allow users to provide and view annotations on custom genome assemblies not hosted by UCSC. Data added to the Genome Browser through these methods are fully integrated with the other tools that UCSC offers, such as the Table Browser, Data Integrator, and the Variant Annotation Integrator.

### Other tools

The UCSC Genome Browser provides numerous command-line and web-based tools for accessing and analyzing data outside of the primary visualization. Interactive tools such as the Table Browser and the Data Integrator allow users to retrieve data from the Genome Browser’s underlying tables. The Table Browser allows users to intersect, filter and download output from these tables in a variety of formats including a BED file or as a custom track loaded directly into the Browser. Output from the Table Browser can also be directly uploaded to external analysis platforms including GREAT (25), Galaxy (26), or GenomeSpace (http://www.genomespace.org). The Data Integrator allows users to combine and extract data from multiple tracks simultaneously (for more details see the “Data Integrator” section below).

The Variant Annotation Integrator (VAI) enables users to obtain the predicted functional effects on different types of annotation tracks from a defined list of variants. Users can upload their variants to the VAI in VCF or pgSnp format or provide a list of dbSNP variant identifiers. The VAI can be used to report the functional effect of a particular variant on a gene prediction, ENCODE regulatory region, or conserved element from the Genome Browser’s Conservation data track. Users can view the output of the VAI in a web browser, or download a file of tab-separated text in Ensembl’s Variant Effect Predictor (VEP) format (27).

Other web-based tools we provide include the Gene Sorter (28), Genome Graphs, In-Silico PCR and VisiGene. The popular BLAT tool can be used to align DNA, RNA, or protein sequences to a genome; another popular tool, LiftOver, can be used to map genomic positions in one genome to another via sequence homology. Both are available on the UCSC Genome Browser website or as downloadable command-line executable files.

We also provide many command-line utilities for extracting useful information from large data files, converting files between related formats, and creating and maintaining data hubs. Prebuilt versions of these tools and their source code are available for Mac OSX and Linux systems (http://hgdownload.soe.ucsc.edu/downloads.html#utilities_downloads). The source code for the Genome Browser is available free for non-commercial or academic use from our secure storefront (https://genome-store.ucsc.edu/) or though git (http://genome.ucsc.edu/admin/git.html). The Genome Browser in a Box (GBiB) (29), which is a virtual machine image of the entire Genome Browser that can be used on a personal computer, is also available for free for noncommercial or academic use from the Genome Browser store. Commercial licenses for the Genome Browser source code and GBiB are also available for purchase from the store.

## NEW AND UPDATED DATA IN THE GENOME BROWSER

### New and updated genome assemblies

Over the last year, we have released a genome browser for one newly sequenced organism and updated browsers for three others (Table 1). These releases include two primate genome browsers: the initial release for the endangered bonobo *(Pan paniscus),* a close relative of chimpanzees and humans, and an update for the tarsier *(Tarsius syrichta),* a small Southeast Asian primate. The genome browsers for two important model organisms, zebrafish *(Danio rerio)* and fruit fly *(Drosophila melanogaster),* were updated as well. Both of these organisms have served as important models for genetics and developmental biology. This release of the *D. melanogaster* assembly marks the first update to that genome in over eight years (30). A new hub containing the genome of a Hawaiian *C. elegans* strain, CB4856 (31), produced by the Waterston Lab at the University of Washington, was added to our list of publicly available hubs.

**Table 1.** New and updated genome browsers

| Common name | Scientific name | Sequencing center | UCSC ID | Seq. ctr. ID |
| --- | --- | --- | --- | --- |
| *New species* |  |  |  |  |
| Bonobo | *Pan paniscus* | Max Planck Institute for Evolutionary Anthropology | panPan1 | Max-Planck Institute panpan1 |
| *Updated species* |  |  |  |  |
| Fruit fly | *Drosophila melanogaster* | FlyBase Consortium, Berkeley | dm6 | BDGP Release 6 + ISO1 MT |
|  |  | Drosophila<br>Genome Project,<br>Celera<br>Genomics |  |  |
| Tarsier | *Tarsius syrichta* | Washington<br>University | tarSyr2 | Tarsius_syrichta-2.0.1 |
| Zebrafish | *Danio rerio* | Genome<br>Reference<br>Consortium | danRer10 | GRCz10 |

### New and updated genomic annotations

In the past year, we have added over 200 new and updated tracks to the UCSC Genome Browser database for our existing assemblies (Table 2), as well as 7 public track and assembly hubs (Table 3). This section highlights some of the major data sets released.

**Table 2.**
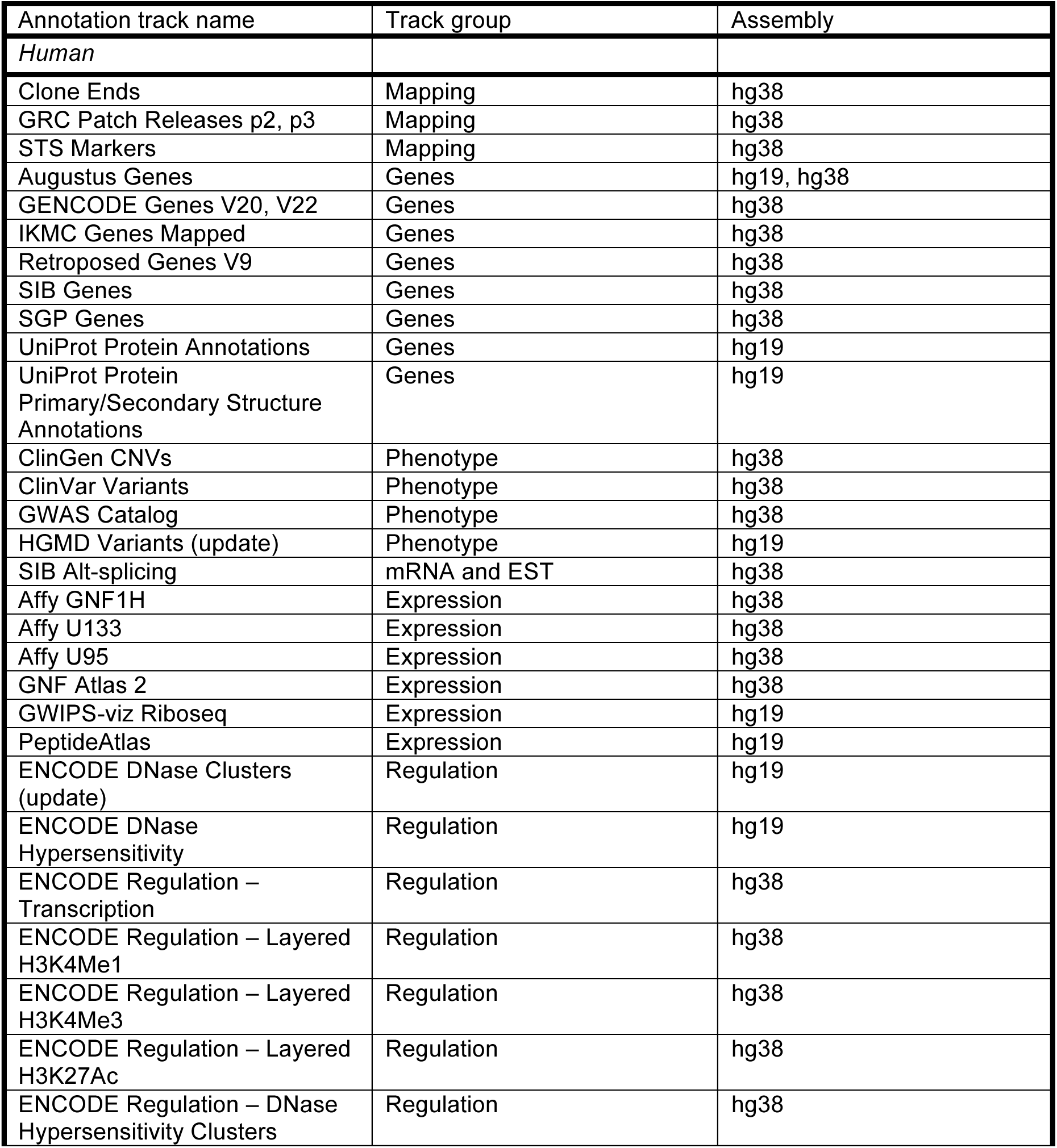
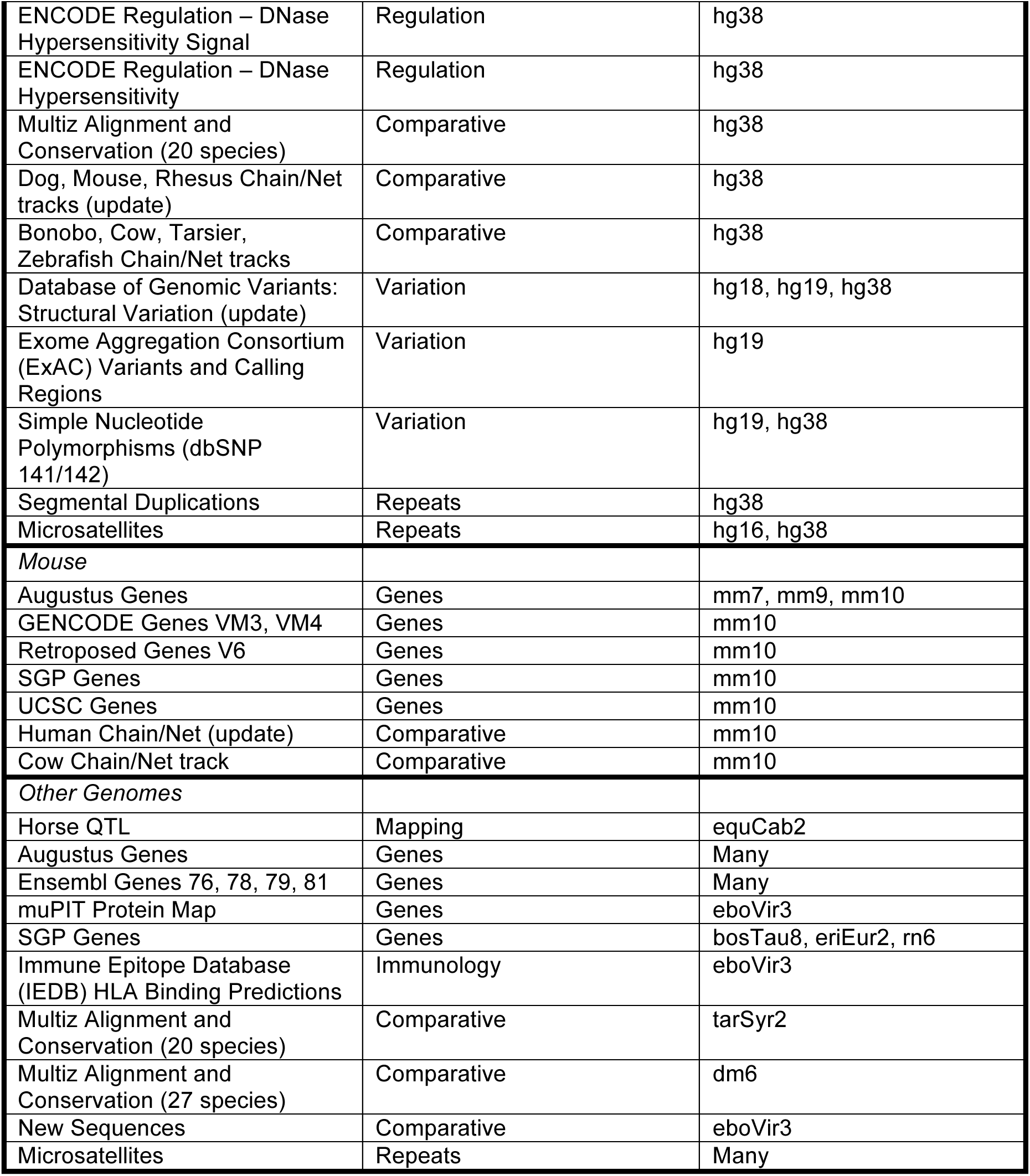
New and updated annotation tracks in the Genome Browser

**Table 3.** New track and assembly data hubs

| Hub Name | Source | Assemblies |
| --- | --- | --- |
| *Track Hubs* |  |  |
| CPTAC Hub v1 | Clinical Proteomic Tumor Analysis Consortium (CPTAC) | hg19 |
| LIBD Human DLPFC Development | Lieber Institute for Brain Development (LIBD) | hg19 |
| PhyloCSF | Broad Institute | hg19, hg38, mm10 |
| Porcine DNA methylation and gene transcription | Laboratory of Comparative Genomics, University of Illinois and Animal Breeding and Genomics Centre, Wageningen University | susSCr3 |
| rfam12_ncRNA | Rfam at the Wellcome Trust Sanger Institute | hg38, mm10, galGal4, danRer7, ci2, dm6, ce10, sacCer3 |
| Vista Enhancers | VISTA Enhancer Group, Lawrence Berkeley Laboratories | hg19, mm9, mm10 |
| **Assembly Hubs** |  |  |
| C_elegans_isolates | Bob Waterston Lab, University of Washington | A C. elegans environmental isolate genome |

#### Gene sets

***UCSC genes.*** The UCSC Genes set contains predicted protein-coding and non-coding genes and their transcripts based on evidence from a variety of sources (32). The pipeline developed by UCSC to generate this gene set incorporates data from RefSeq, GenBank, CCDS (33), Rfam (34), and gtRNAdb (35) to build a high confidence set of gene and transcript predictions for our two primary assemblies, human and mouse. We have updated the UCSC Genes track available on the default mouse assembly, mm10/GRCm38. This update includes 1,602 brand-new transcripts, increasing the total number to 63,244 transcripts. The total number of canonical genes in the UCSC Genes set increased from 32,408 to 32,958.

Beginning with the hg38/GRCh38 assembly, we have discontinued further releases of the UCSC Genes on the human genome. In an attempt to help the bioinformatics community shift towards a common, high-quality gene set, we have instead decided to use the GENCODE Genes set (below) as the default gene annotation for human assemblies.

***GENCODE genes.*** The GENCODE Project provides a set of pseudo-, protein-coding, and noncoding RNA genes for both the human and mouse genomes. The GENCODE Genes set combines predicted transcripts from Ensembl’s genebuild pipeline and manually curated transcripts from HAVANA to create a high-confidence set of gene and transcript annotations. We have updated the track on the mm10/GRCm38 mouse assembly twice in the last year: first to VM3 and then most recently to VM4. The VM4 release covers 43,346 genes with a total of 103,639 transcripts. The track on the hg38/GRCh38 human assembly has been updated twice, first to V20 and then to V22, which has replaced UCSC Genes as the default gene set on the human genome. The Genome Browser keeps the older versions to allow continuity for those using an older version. The V22 release (Figure 1) covers 60,498 genes with a total of 198,619 transcripts, which represents a 2% increase in the number of transcripts from V20. A page of detailed information that replicates the detail formerly displayed on the UCSC Genes track now accompanies each transcript in the GENCODE Genes track. This includes imputed haplotype allele statistics based on the 1000 Genomes Project (36), microarray expression data from GNF Atlas (37), protein domains from InterPro (38) and Pfam (39), three-dimensional protein structures from PDB (40), and biological pathway interactions from KEGG (41) and Reactome (42, 43). We also make an effort to include links to more information on other resources, such as BioGPS (44), UniProtKB (45), and Ensembl, for each gene when available.

**Figure 1.**
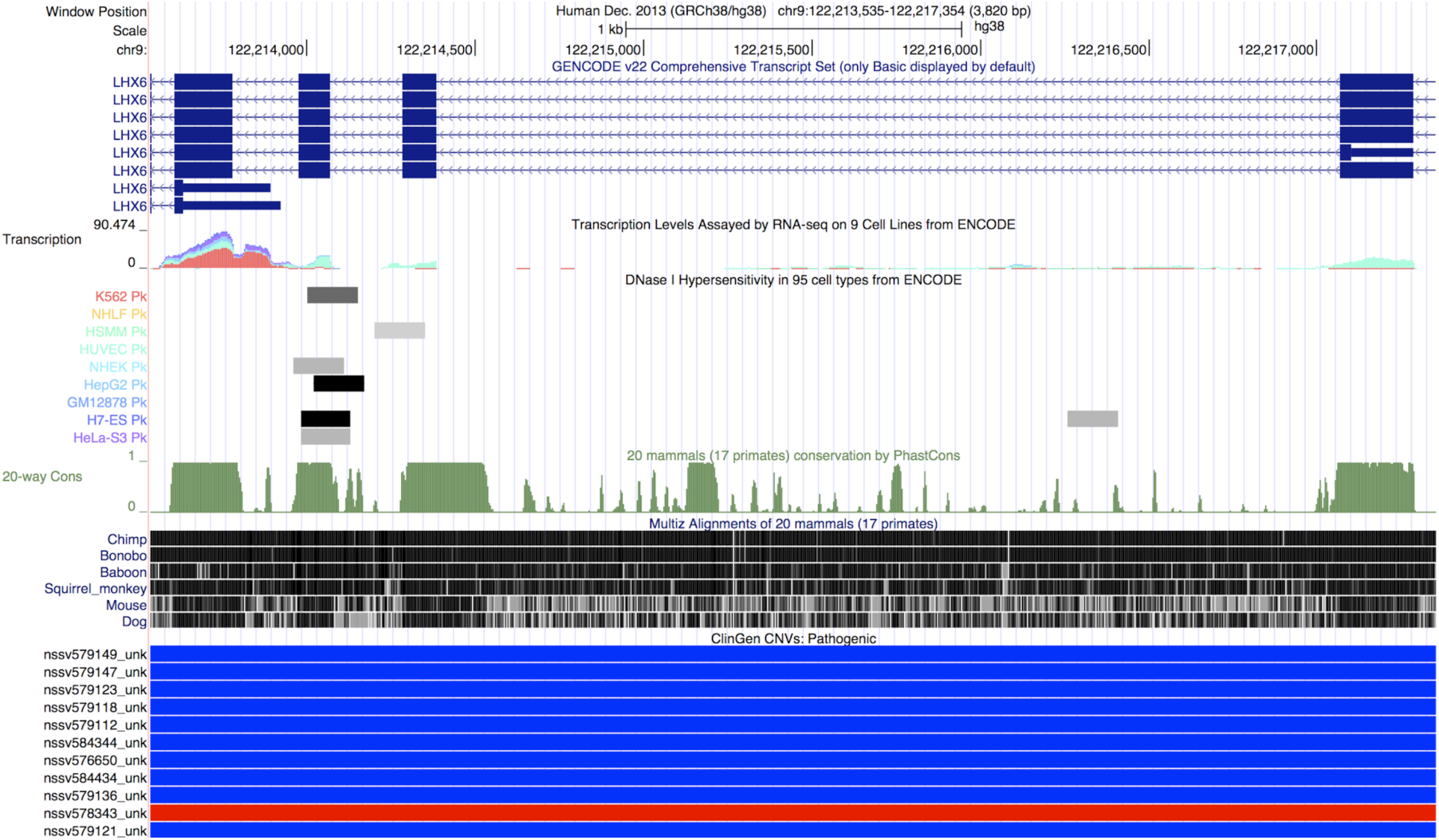
Genome Browser screenshot focused on a region of the LHX6 gene that highlights a selection of the new tracks added in the previous year for the hg38/GRCh38 human assembly. The tracks shown in this display (from top to bottom) include GENCODE Genes V22, transcription levels assayed across 9 ENCODE cell lines, DNase hypersensitive regions based on data from 95 ENCODE cell lines, genome-wide conservation scores calculated using phastCons, a multiple genome alignment created using Lastz and Multiz, and pathogenic CNVs from the ClinGen database. This screenshot demonstrates how one can use the ENCODE transcription levels and DNase hypersensitive regions in the Genome Browser to identify expression of different transcripts in different cells. The multiple alignment and conservation scores can be used to look for other potential regions of importance. Lastly, one can use the ClinGen CNVs and other pathogenicity data in the browser to determine what role, if any, a particular gene plays in disease.

***Augustus genes.*** The ab-initio gene-finding program Augustus (46, 47) uses statistical modeling of gene features to find and annotate potential protein-coding genes, including their transcript splice variants and UTRs. We worked with the program’s creator, Mario Stanke, to create an initial set of Augustus predictions on more than 150 assemblies. We plan to make the Augustus gene prediction track a standard annotation on every new genome browser released at UCSC.

*Phenotype and variation.* Several new clinical variation tracks have been released on hg38/GRCh38. The ClinVar track displays detailed information on simple nucleotide polymorphisms (SNPs) and copy number variants (CNVs) from NCBI’s ClinVar database. ClinVar is a freely available, public database that links variants to their phenotypic characteristics along with supporting evidence (48). The GWAS Catalog track displays a collection of over 100,000 SNPs curated by EMBL-EBI (previously by NHGRI) and collected from a number of published genome-wide association studies (49). The ClinGen tracks (Figure 1) display CNVs identified in patients who have undergone genetic testing for various developmental disorders. ClinGen is a public database of clinical variants that aims to collect and share data among clinicians, patients and researchers and provide them with information on a variant’s clinical relevancy and pathogenicity (50). All three of these tracks are updated in the Genome Browser on a regular basis to reflect the newest data from their sources.

We have added two tracks containing data from the Exome Aggregation Consortium (ExAC) (http://exac.broadinstitute.org) for hg19/GRCh37. The consortium aims to collect and bring together exome sequencing data from a number of large-scale and varied exome sequencing projects. It currently contains 60,706 exomes and their variants collected from its member projects. These tracks contain the ExAC calling regions and over 8.75 million single nucleotide variants (SNVs) in these calling regions collected from the various contributors.

We loaded two releases of dbSNP, v141 and v142, into the Genome Browser database this year, which include millions of variants across both hg19/GRCh37 and hg38/GRCh38. The v142 release includes over 50 million new variants from the 1000 Genomes project Phase 3. We plan to release a set of tracks for dbSNP v144 by the end of 2015.

*Gene Expression.* We have added new microarray expression tracks based on data from GNF Atlas 2 and the Affy probe sets GNF1H, U133, and U95 to hg38/GRCh38. Two sets of proteomics data from PeptideAtlas and the Clinical Proteomic Tumor Analysis Consortium (CPTAC) are now available for hg19/GRCh37. The data from PeptideAtlas are based on over 971 samples collected from laboratories around the world and identifies over 1,021,823 distinct peptides covering 15,136 proteins (51, 52). The CPTAC data are available as a public track hub, and cover mass spectrometry data across breast, colon, and ovarian cancer samples from The Cancer Genome Atlas (TCGA) (53).

*ENCODE regulatory data.* We have been migrating ENCODE regulatory data from hg19/GRCh37 to hg38/GRCh38. We recreated the DNase hypersensitivity signals (Figure 1) and hotspots across 95 cell lines from the raw source files using the UCSC ENCODE DNase analysis pipeline on hg38/GRCh38. We “lifted” (a process that involves mapping coordinates from the previous assembly to the newer one) transcription levels across nine ENCODE cells lines (Figure 1) and histone methylation and acylation marks from seven cell lines from hg19/GRCh37 to hg38/GRCh38.

*Comparative genomics.* We have added numerous conservation and pairwise alignment tracks over the past year. Conservation tracks at UCSC contain a multiple alignment constructed using Lastz and Multiz, conservation scores across the genome calculated using both phastCons and phyloP, and conserved regions identified with phastCons. Newly released conservation tracks this year include 20-way alignments on human (hg38/GRCh38) (Figure 1) and tarsier (tarSyr2) that include 17 primate genomes and 3 other mammals, and a 27-way alignment on *D. melanogaster* (dm6) that is primarily focused on 23 species from the *Drosophila* genus, but includes 4 other insect species as out-groups.

## NEW SOFTWARE FEATURES IN THE GENOME BROWSER

### Data Integrator

The Data Integrator (http://genome.ucsc.edu/cgi-bin/hgIntegrator) is a powerful, easy-to-use tool for quickly combining and exporting data from multiple tracks simultaneously. Users can combine up to five tracks from a single genome assembly, selecting from tracks hosted by UCSC or those uploaded to the browser using custom tracks or track data hubs. The Data Integrator will output data from the selected tracks based on their intersection with a primary track, which can be changed at any time by rearranging them. By default, all of the fields from the selected tracks will be included in the output, but this can be reduced to a user-selected subset of fields if desired.

The Data Integrator is available from the “Tools” menu. For more information on the Data Integrator, see http://genome.ucsc.edu/goldenPath/help/hgIntegratorHelp.html (accessible from the “Help” menu on the Data Integrator web page)

### Enhancements to data hubs

*BLAT your genomes.* Users are now able to specify a remote BLAT instance for use with an assembly data hub. After proper configuration, users of a hub can then use the web interface of BLAT to align sequences with their genome.

*More robust gene prediction display.* Both track and assembly data hubs now support the use of the bigGenePred file format. The “genePred” format is a table format used at UCSC for the display of gene prediction tracks such as UCSC Genes or Ensembl Genes. This format supports a more robust display of gene predictions with features such as codon translation and exon numbering. Similar to other UCSC “big” formats such as bigWig and bigBed, bigGenePred is a compressed, binary format and can be produced using command-line tools provided by UCSC and loaded into the Genome Browser. Information on creating a bigGenePred track can be found at http://genome.ucsc.edu/goldenPath/help/bigGenePred.html.

*Use your data hubs across genome browsers.* As our track and assembly data hub features have grown into widely used methods for organizing and sharing genomic data throughout the research community, there has been increasing demand to use these data hubs across a variety of genome browsers hosted at different sites, including Ensembl, Biodalliance (54), and the Epigenome Browser at Washington University in St. Louis (55). To facilitate this, we are collaborating with these groups to version the “trackDb” configuration settings that are used in data hubs to define attributes of a track (such as the track label, data type, and default display mode, among other things) and to define support levels for subsets of these settings. As part of this effort, all of our current settings have been assigned a support level: “required” (those that a trackDb must include), “base” (common settings supported at other sites), or “full” (all other settings). The current trackDb setting specification, which has been released as version ‘v1’, is viewable at http://genome.ucsc.edu/goldenPath/help/trackDb/trackDbHub.html. We plan to increment the version as new settings are added. These additions will allow engineers of other Genome Browsers to declare which trackDb versions and levels they support, thus enabling users to determine if their hubs will display correctly on a given browser.

We have incorporated these new versioning and support level features into our existing hubCheck tool, which is used to validate and test the integrity of a data hub. The tool now allows users to validate their data hubs against these new trackDb support levels and version numbers or, alternatively, against a custom set of trackDb settings.

### Clinical variants beacon

In collaboration with the Global Alliance for Genomics and Health (GA4GH) (http://genomicsandhealth.org/), we have created the new Clinical Variants Beacon (http://genome.ucsc.edu/cgi-bin/hgBeacon), which allows users to programmatically query two of the clinical variation tracks available in the UCSC Genome Browser: the Leiden Open Variation Database (LOVD) (56) and Human Gene Mutation Database (HGMD, public version) (57). Users can submit URL queries to the beacon that contain the position, the data set (LOVD or HGMD), the reference genome, and the alternate base. It will return plain text or a JSON-formatted string indicating whether the alternate base is found in the specified database at that position. Users can then navigate to the source database to learn more about the allele at that position.

### Improvements to the browser display

*Exon-number display.* You can now quickly find the exon number of a transcript by hovering your mouse cursor over the exon in the browser display. This feature will display the chosen exon out of the total number of exons in a particular transcript (e.g. “Exon 6/8”). This feature also works with introns.

*Display data on both strands.* Many users find it convenient to display continuous “wiggle” graph data, such as transcription levels, on the reverse strand as negative numbers. Users can now use the “negate values” option to quickly negate the values in the wiggle file (positive values become negative and vice versa).

### Expanded training and workshop program

We have expanded the online training offered by the UCSC Genome Browser. Over the past year, we have created a series of short videos that demonstrate stepwise solutions to common questions such as obtaining a list of genes in a specified region or finding the SNPs that occur within a particular gene. All of the videos are available through YouTube and are accompanied by a full transcript of the voice-over. A full list of videos and links to their transcripts is available here: http://genome.ucsc.edu/training/vids/index.html.

We also continue to update and create new training documentation for the various Genome Browser tools. Some of the most exciting additions this year include the Genome Browser blog (http://genome.ucsc.edu/blog/) and a number of track and assembly data hub “Quick Start” guides. Our blog allows us to give in-depth looks at new features, new data sets or other interesting topics related to the Genome Browser. The Quick Start guides provide an easy-to-follow step-by-step process that one can use when creating a data hub, as well as example files and commands to copy these examples to a local computer for manipulation and testing.

We have also expanded the number of on-site workshops we offer. These workshops are tailored to the experience level of each group and can cover topics ranging from the Genome Browser basics to complicated multi-step queries across the different Genome Browser tools. Over the last year, our training team has given over 35 presentations and training sessions around the world ranging from small rooms of 15 people to conference halls seating hundreds. To inquire about hosting a workshop at your institution, see: http://bit.ly/ucscTraining.

## FUTURE PLANS

We have plans to continue adding and updating vertebrate genome assemblies as they become available in NCBI’s GenBank repository, and will continue to update and release new annotations for the various assemblies hosted at UCSC. We intend to focus our efforts on working with data contributors to expand our annotation collection on the hg38/GRCh38 assembly with both new data sets and popular data sets migrated from previous assembly versions. These include annotations across all of the major data types including gene predictions, phenotype and disease variants, regulatory regions and expression level data sets. Along with these annotations, we plan to make hg38/GRCh38 the default human assembly in the Genome Browser by late 2015. This means that users will be automatically directed to hg38/GRCh38 when accessing the UCSC Genome Browser for the first time. Older assemblies of the human genome will continue to be accessible.

One of the major data sets that we plan to release in the coming year is based on data from the Genotype-Tissue Expression (GTEx) project (58). This project contains data aimed at aiding researchers in identifying variation in expression levels between different tissue types and then relating these variations back to genomic variation. The GTEx data will also include new visualization modes that will highlight differences in expression between tissues and allow researchers to relate these differences back to genotype.

We are developing a new “multi-region” mode that allows users to visualize more than one region at a time from a single assembly in the same window. We envision this to have three primary uses: (1) an exon-only mode that excludes intronic sequence from the browser display, (2) a means for viewing haplotype sequences in the context of the reference genome, and (3) a mode that allows for user-defined regions. Other major software features planned for the coming year include the ability to generate density plots for BAM and other track types on the fly, and improvements and additions to the Data Integrator tool.

## CONTACT US

The UCSC Genome Browser maintains a number of actively monitored mailing lists and social media channels. Questions about the Genome Browser and the resources we provide can be directed to our publicly archived, searchable mailing list. Questions involving sensitive information can be sent to the private mailing list. To receive updates about Genome Browser news including the release of major data sets, new assemblies, and new software features, subscribe to the mailing list or follow us on Twitter (@GenomeBrowser). For more contact information, see our Contacts page at http://genome.ucsc.edu/contacts.html.

## ACKNOWLEDGEMENTS

We would like to thank our many data contributors and collaborators whose work make the Genome Browser what it is, our loyal users for their support and vital feedback, and our exceptional group of system administrators who keep the Genome Browser running.

## FUNDING

This work was supported by National Human Genome Research Institute [5U41HG002371 to G.P.B., J.C., H.C., C.E., P.A.F., L.G., M.H., S.H., A.S.H., D.K., W.J.K., R.M.K., K.L., B.T.L., B.P., B.J.R., K.R.R., M.L.S., A.S.Z.; 5U41HG006992 to W.J.K., K.L., B.T.L., B.J.R; 5U41HG007234 to M.D., R.A.H., B.P.; 5R25HG006836 to P.N.]; National Cancer Institute [5U54HG007990 to M.D., M.H., R.M.K., B.P.; 5U24CA180951 to M.D.]; California Institute for Regenerative Medicine [GC1R-06673-C to G.P.B., L.G., M.H., S.H., D.K., W.J.K., B.T.L., K.R.R., A.S.Z.]; Howard Hughes Medical Institute [090100 to D.H.]; Simons Foundation [351901 to B.P.]; and the UCSC Jack Baskin School of Engineering Dean’s Office Fellowship [to P.N.]. Funding for open access charge: National Human Genome Research Institute.

*Conflict of interest statement.* G.P.B., H.C., J.C., M.D., P.A.F., L.G., D.H., M.H., R.A.H., S.H., A.S.H., D.K., W.J.K., R.M.K., B.T.L., K.L., B.J.R., K.R.R., M.L.S. and A.S.Z. receive royalties from the sale of UCSC Genome Browser source code and GBiB licenses to commercial entities. W.J.K. works for Kent Informatics.

